# Yellowhammer (*Emberiza citrinella*) males sing using individual isochronous rhythms and maximise rhythmic dissimilarity with neighbours

**DOI:** 10.1101/2025.06.17.660106

**Authors:** Anna N Osiecka, Marcos Q Oliva, Jan Kouřil, Tereza Petrusková, Lara S Burchardt

**Affiliations:** Institute for Theorethical Biology, Humboldt University of Berlin, Berlin, Germany; ENES Bioacoustics Research Laboratory, University of Saint-Etienne, Saint-Etienne, France; Escuela de Biología, Universidad de Costa Rica, San José, Costa Rica; Department of Ecology, Faculty of Science, Charles University, Prague, Czechia

**Keywords:** birdsong, individuality, isochrony, rhythm adjustments, rhythmic patterns, vocal communication

## Abstract

The temporal structure of an animal’s vocal output can be cognitively controlled, presenting an interesting aspect for vocal learning species that benefit from diversification. Although birdsong is the most thoroughly studied aspect of animal communication, its rhythms remain largely unknown. Here, we revisit the question of vocal individuality in the songs of male yellowhammers (*Emberiza citrinella*) purely from a rhythmic perspective. Yellowhammers use simple songs composed of two phrases: the *initial phrase*, with one individual usually having a repertoire of on average two such phrase types, and the *dialect* phrase containing the dialect that is the same for all males at a given locality. Some of the initial phrases are commonly shared between various males, but their combinations and frequency contours are individually unique and temporally stable. Using focal recordings of 38 known individuals, collected over three years in the same geographic location, we calculated a set of nine temporal indices to describe each song and compared individuals in a permuted discriminant function analysis. Subsequently, we calculated the potential for individual coding for each of these parameters. To assess whether rhythmic similarity may depend on the singers’ proximity, we calculated vocal dissimilarity as the Euclidean distances between each two males, and used Kendall’s correlation. We show that yellowhammer males use individual rhythms, maintained over different phrase types and carried mostly in the inter-onset-interval variability and syllable rate, as in syllables per second, within the song. There were particularly strong rhythmic differences between singers among the closest neighbours, which decreased over the first 600 m. There was no pattern for neighbours beyond this distance. This study is the first to demonstrate the existence of strong individuality based purely on rhythm, as well as rhythmic differentiation from neighbours, in a songbird.

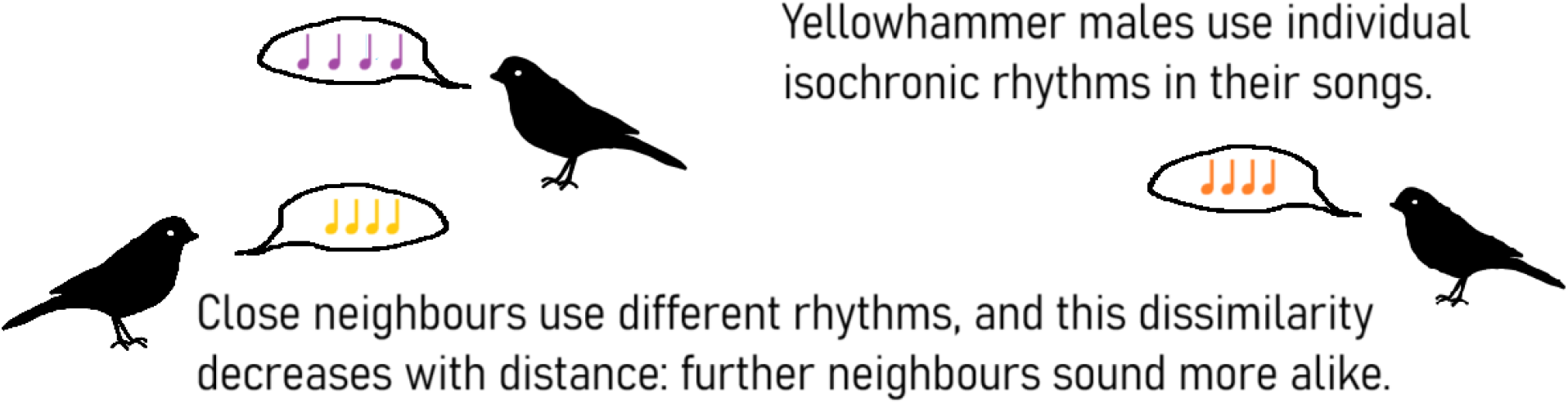

## Introduction

Avian vocalisations tend to vary at geographical, population, and individual scales. These are also the scales at which this variation is usually considered. In the studies of vocal individuality, we tend to search for a kind of “global individuality” - the overall differences between birds in a given population. At the same time, we know that social bonds can influence the vocal output of an animal, making them more similar to their social group (e.g. the African penguin, *Spheniscus* demersus Baciadonna et al. (2022)) or partners (e.g. ravens Corvus corax Luef et al. (2017), and little auks Alle alle Osiecka et al. (2023)). It seems likely that other social factors may influence vocal similarity between animals - for example, some animals might benefit from maximising recognisability from their neighbours, especially if territories or leks are kept **?**, and that individuality exists rather in relation to the others.

Vocal individuality is usually assessed on the basis of the spectral parameters of an acoustic signal or its frequency contour (similarly to a vocal signature). Yet the temporal structure of vocalisations can prove at least equally important to identity encoding Mathevon et al. (2017) Robisson (1992) Osiecka et al. (2024) and perception of vocalisations Ter-Mikaelian et al. (2013). This can prove particularly interesting in species with elaborate and cognitively modified vocal behaviour, and in territorial species benefiting from being easily recognised by the others. Songbirds are a very good example of such species, and moreover, they are known to discriminate among individuals very well. However, the cues they use to reveal singer’s identity remain unclear. When you think of birdsong, you likely imagine a melody, or rhythm Bilger et al. (2021).

### Can cues to singer identity be coded by the rhythms of their songs

The yellowhammer *(Emberiza citrinella*) is a common passerine. During the breeding seasons (May-August), yellowhammer males establish territories of up to approximately 1 hectares (circa 150 m in diagonal Møller (1990)), located along hedges or woodland borders, from where they sing songs ascribed largely to territorial behaviour Hiett & Catchpole (1982). Yellowhammer song is composed of a series of single- or multi-element syllables composing the *initial phrase*, followed by typically two longer syllables, referred to as *dialect phrase*. While the dialects have been examined for over a hundred years Petrusková et al. (2015), studies on initial phrases are much scarcer, although it is known that their combinations are individually unique and temporally stable Hansen & Balsby (2012). At the same time, initial phrases can be shared across individuals Rutkowska-Guz & Osiejuk (2004), Hansen & Balsby (2012) (Fig. 1). The *initial phrases* are composed of syllables produced in roughly isochronous sequences, making them a great candidate to study individual traits in simple rhythms.

**Figure 1:**
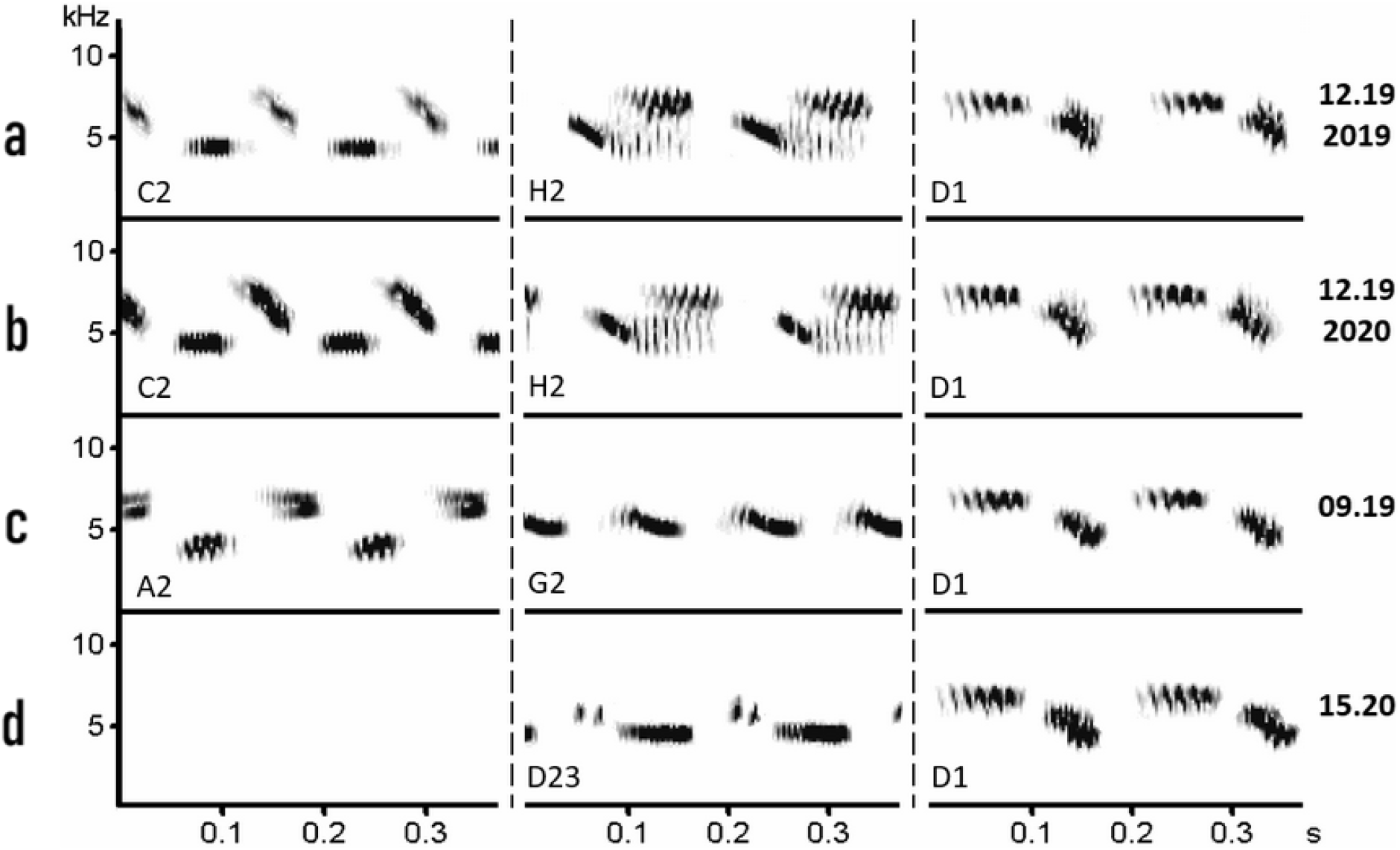
Examples of initial phrase repertoires of three different males Panels a and b show the same male in different years. Panels c and d show two different males. Note that phrase D is shared by these three males. While the phrase type syllable repertoire (columns) used by yellowhammer males (rows) is to some extent shared by individuals, their syllable contours remain individually distinctive.

Here, we revisit an existing library of male yellowhammer songs to look at their individuality from a purely rhythmic perspective on a small geographical scale. We investigate whether individual singers may differ in the rhythmic structure of their songs, and whether these differences might depend on the distance between the singers within a single location.

## Methods

### Ethical note

This study is based on previously existing data, and no specific permits were necessary. The previous work that provided these data adhered to all applicable international, national, and institutional guidelines for research on animals. Data were collected using a non-invasive recording technique from a distance, and no birds were captured.

### Location and fieldwork

Yellowhammers were recorded at the foothills of the Brdy mountain range in central Bohemia, Czech Republic, from 2019 to 2021. The study area (49.844N, 14.103E; roughly 2 km in diagonal) provides optimal habitats for this species - meadows and fields interspersed by belts of bushes and trees Cramp & Perrins (1994).

The area was visited at least once every two weeks during the breeding season of Yellowhammers (May to August Cramp & Perrins (1994)), and all encountered singing males were recorded with a directional microphone (Sennheiser ME 67) and a digital recorder (Marantz PMD661; 16 bit, 44.1 kHz sampling rate). Most of the recordings were collected by JK, a minority by TP. For each recording, we noted the date, time, location, and behaviour of the male. If more males were recorded at the same time (i.e. in case of singing interaction, counter-singing), they were marked as “male 1”, “male 2” and so further directly as a voice comment in the recordings. To avoid misidentifications of the males, only recordings with songs of a single male (i.e. only solo, and not counter-singing) were used in subsequent analysis.

### Data availability

Full data and scripts generated in this work, as well as supplementary materials are available at: https://osf.io/jwt3g/?view_only=a971401f45744a709bb8accd0ee9769f

### Song analysis

*Identification of males and recording pre-processing* All recordings were first visually inspected by JK in Avisoft SASLab Pro Specht (2002) as sonograms (FFT Length = 256, Frame % = 100, Window = Hamming). Based on this visual inspection, song phrases were assigned to distinct phrase types based on their temporal, structural and frequency characteristics, and phrase repertoires were determined for each recording. Recordings with matching phrase repertoires were then assigned to individual males (following the methods of Petrusková et al. (2016)). The syllable types used per individual are shown in Supplementary Table 1.

**Table 1.** Definitions of the temporal parameters calculated for each song. Note that here and throughout the article, the term *song* only refers to the *initial phrase*.

| Parameter | Definition |
| --- | --- |
| Mean syllable duration (s) | Mean duration of individual syllables in a song |
| Mean IOI (s) | Mean inter-onset-interval between each two consecutive syllables |
| CV syllable | Coefficient of variation of syllable duration in a song, calculated as $\frac{SD_{\text{syl\_dur}}}{\bar{x}_{\text{syl\_dur}}} \times 100$ |
| CV IOI | Coefficient of variation of inter-onset-intervals in a song, calculated as $\frac{SD_{\text{IOI}}}{\bar{x}_{\text{IOI}}} \times 100$ |
| Duration (s) | Duration of the phrase part of the song, i.e., excluding dialect syllables (if present) |
| No. syllables | Number of phrase type syllables in a song |
| Syllable rate (Hz) | Number of syllables per second |
| Rk global min | The minimum value of the song's integer ratio, calculated for all syllable pairs in the song, Equation 1 |
| Rk global max | The maximum value of the song's integer ratio for all syllable pairs in the song, Equation 1 |

Recordings were manually annotated in Raven Pro 1.6 (Cornell lab of Ornithology R Core Team (2024)) by two annotators (MQO and ANO), with the following spectrogram settings: Hann window, window size 214, contrast 50, brightness 62, colour scale grey 1st, and bandwidth filter from 1.5 to 15 kHz. The exact beginning and end of each syllable was marked, and the syllable type noted (based on the previous Avisoft classification). A total of 633 songs composed of 52 phrase types and sung by 38 individual males have been annotated for further analysis (see Supplementary Table 1). Five of the males were recorded in two years, and one over three years, all other males were only recorded in one year.

*Rhythmic indices* Only *initial phrase* syllables were used for the calculations (i.e., dialect syllables were not considered). Nine temporal parameters were calculated using a custom-written R script, which was published alongside the project data and is based on Burchardt et al. (2025) Osiecka et al. (2024) (Table 1, Figure 2). Integer ratios, as first described in Roeske et al. (2020), are calculated not only for adjacent intervals in the sequences, but for all interval pairs of a sequence, allowing for a more global interpretation of isochrony Burchardt et al. (2025), Equation 1.

**Figure 2:**
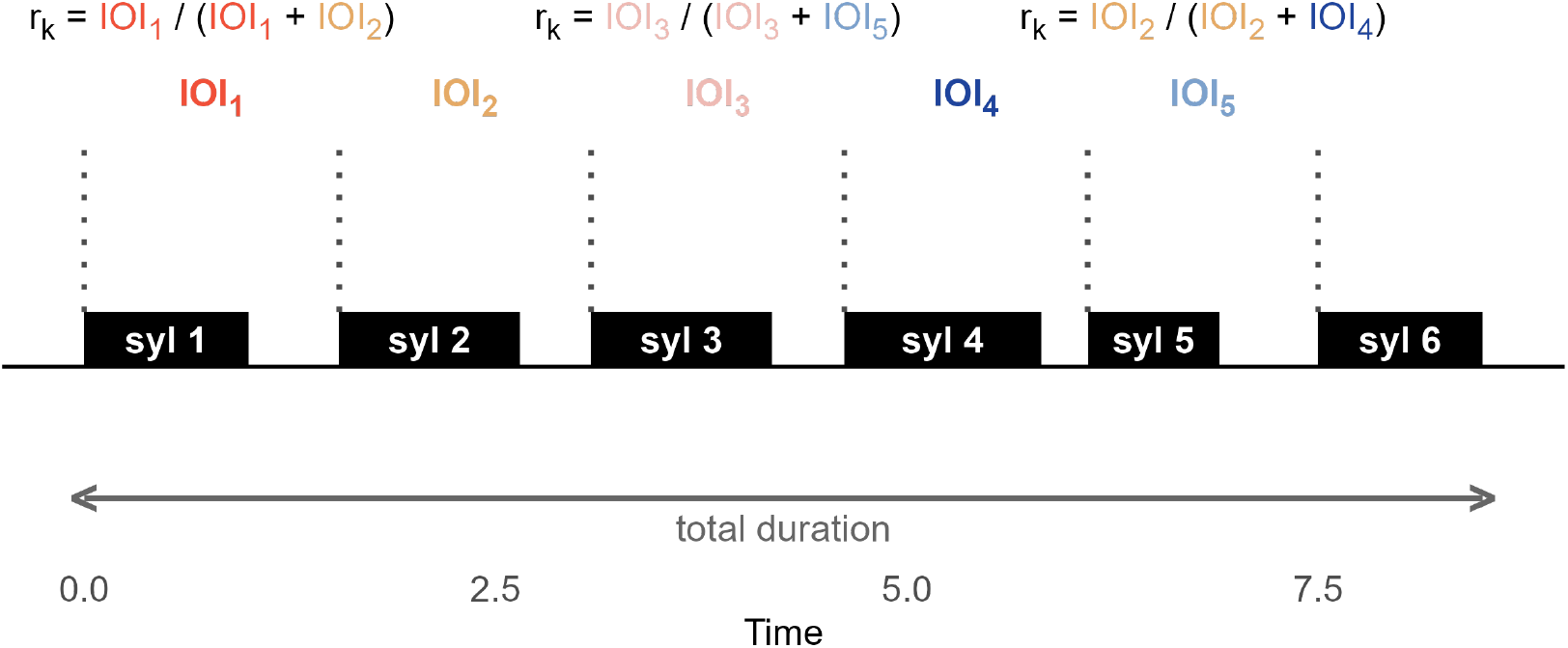
Visual representation of some of the rhythmic parameters based on a random isochronous sequence. See how the global integer ratios (rk) are calculated for any pair of intervals, not just the adjacent ones. Note that the given rk calculations serve as an example, and are not an exhaustive list.

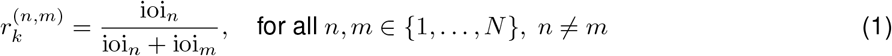

Since the mean and median global integer ratio values were both 0.5, we considered the minimum and maximum global values in subsequent analyses of individuality (see Fig 3 for the distribution of integer ratios).

**Figure 3:**
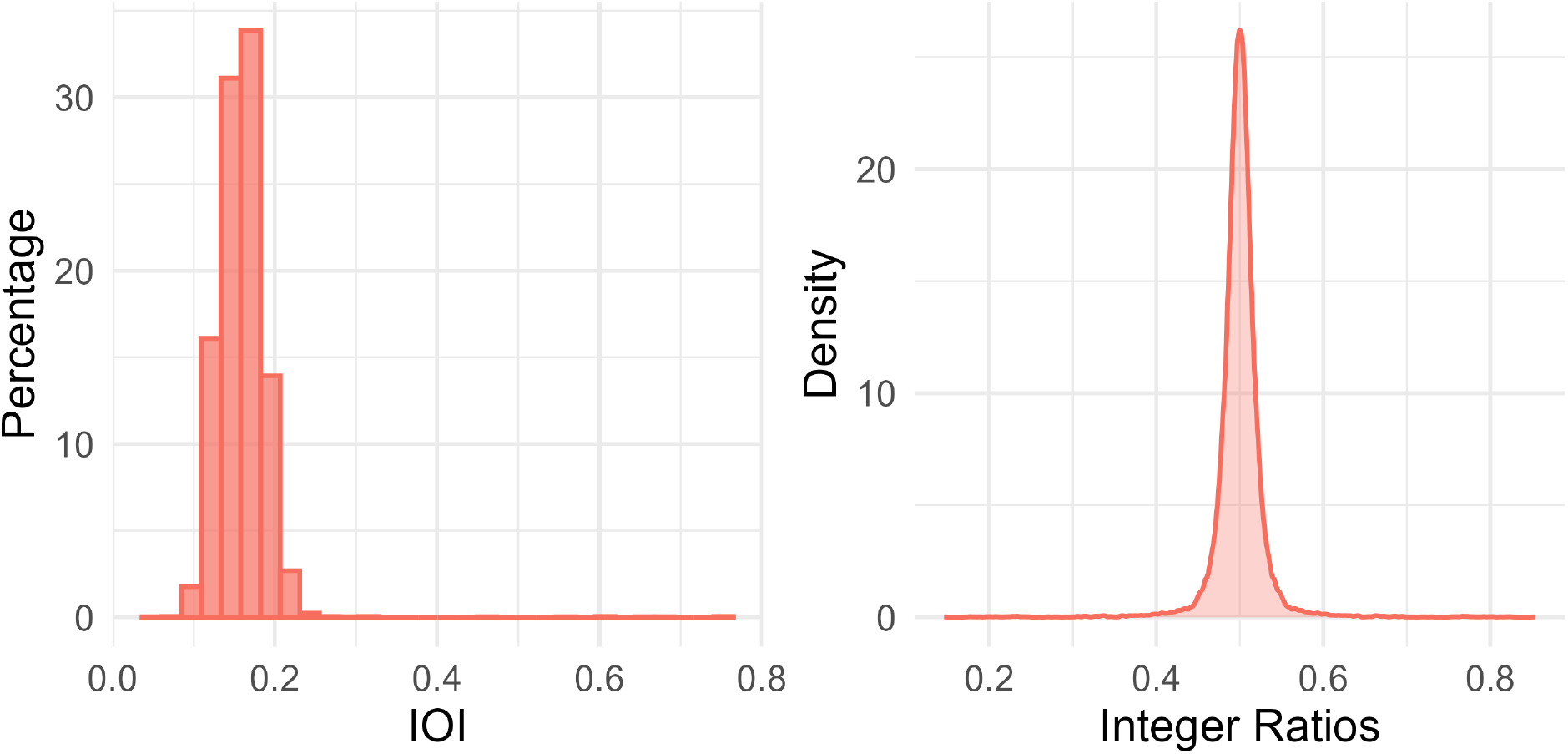
Density plots of the observed IOIs (left panel) and integer ratios (right panel). Both measures indicate a clear isochronous pattern in the yellowhammer song

### Are rhythms individually specific?

The suitability of the data for factor analysis was checked using the KMO function of the EFAtools package Steiner & Grieder (2020). The data were not suitable for PCA. Thus, raw variables were used as input for the permuted discriminant function analysis (pDFA), which allows balancing the uneven sampling of individuals, and controlling for other parameters Mundry & Sommer (2007). We performed a pDFA that pooled all available songs from all individuals, phrase types, and from all years. The pDFA with nested design was performed using the pDFA.nested function of a script provided by Roger Mundry (based on the lda function of the MASS package Ripley et al. (2013)). Two tests were run: (1) using individual ID as a test factor and no restriction factor, to see whether rhythms are individually specific across phrase types, and (2) using individual ID as a test factor and phrase types as a restriction factor, to see whether this remains true within the phrase types. We ran a total of 1000 permutations for each analysis.

To test which of the parameters carried the most information about the singer’s identity, we calculated the Potential of Individual Coding (PIC Robisson (1992)) using the calcPIC function of the IDmeasurer package Linhart (2019).

### Does dissimilarity decrease with distance

*Distance between callers* To assess how distance between callers can affect their similarity, each two birds were paired in dyads. The dataset was first split by year since some individuals were recorded in more than one year. We calculated the distances for each dyad (i.e., the distance between each two callers) using the singers’ coordinates in a distHaversine function, geosphere package Hijmans et al. (2017), which calculates the shortest way between two points on an ellipse.

*Dissimilarity scores* Following Luef et al. (2017), rhythmic dissimilarity was calculated as a Euclidean distance matrix (dist function, stats package R Core Team (2024)) based on the mean values per individual of the nine temporal indices we calculated. The obtained values are further on refered to as the *dissimilarity score*. Dissimilarity scores of 0 indicate perfect similarity, like for self-correlation, the higher the dissimiliarity score, the less similar the dyad.

*Correlation test* The data did not meet the assumptions of a linear model (tested with check_model function of performance package Lüdecke et al. (2021)). We used a non-parametric method, Kendall’s correlation test (cor.test function, stats package R Core Team (2024)), to test the correlation between rhythmic dissimilarity and the distance between each two singers. Kendall’s correlation was used to avoid assumptions of normal data distribution.

Self-correlations were removed from the dataset before fitting the models. We split the data in eight groups: (1) singers up to 50 m distance, i.e. closest neighbours, (2) singers up to 150 m distance, i.e. at approx. one territory size, (3) up to 300 m, (4) up to 450 m, (5) up to 600 m, (6) up to 750 m, (7) up to 900 m, and (8) up to 2000 m, i.e. including all available dyads. This was done to allow us to look for patterns at a behaviorally/communicatively useful distance estimated at one quarter of the total site diagonal or equivalent to multiplications of an estimated territory size diagonal, and (b) overall distance patterns not related to the behaviourally/communicatively useful ranges.

## Results

### Are rhythms individually specific

The songs of male Yellowhammers could be classified according to the correct individual above the chance level (p = 0.001) based solely on the rhythmic indices, both across and within phrase types (Table 2, Figure 4) across the years. However, the low relative cross-classification level (below 1) within phrase types and clear grouping by syllable type (see the clickable 3D UMAP figure in the Supplementary Materials) suggest that some rhythmic patterns correspond to phrase type rather than to the individual.

**Table 2.** Results of the permuted discriminant function analysis classifying each song to the singer across and within phrase types and across years.

| Result | Across phrases | Within phrase type |
| --- | --- | --- |
| No. of syllable types | 52 | 52 |
| No. of individuals | 38 | 38 |
| Total number of songs | 633 | 633 |
| Correctly classified (%) | 44.01 | 43.42 |
| Chance level for classified (%) | 16.45 | 34.53 |
| P value for classified | 0.001*** | 0.004** |
| Correctly cross-classified (%) | 25.85 | 25.89 |
| Chance level for cross-classified (%) | 2.61 | 16.57 |
| Relative cross-classification level | 1.57 | 0.75 |
| P value for cross-classified | 0.001*** | 0.001*** |

**Figure 4:**
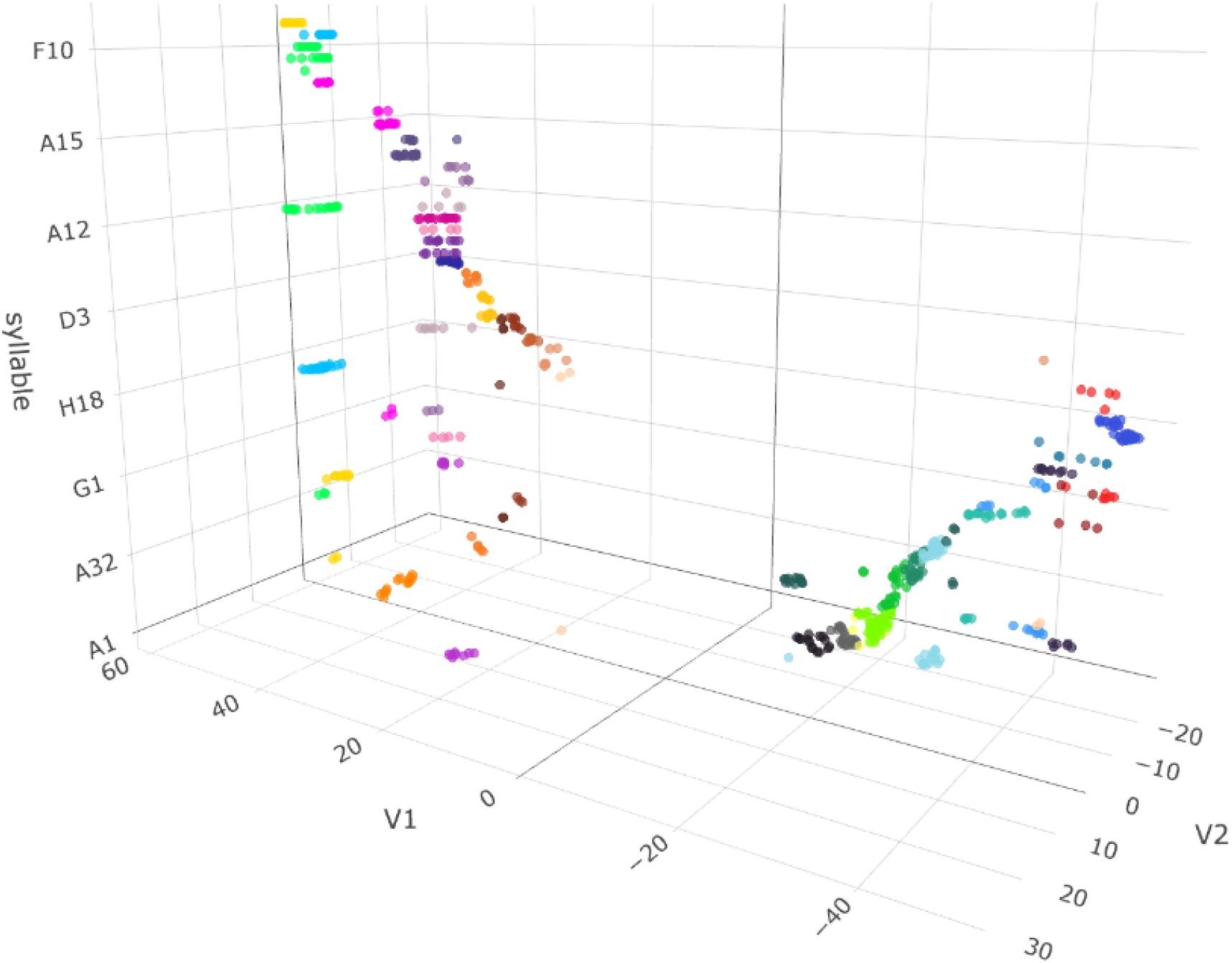
S-UMAP grouping songs based on their rhythmic patterns, prepared using the umap function of uwot package Melville et al. (2022) with the following algorithm settings: 100 neighbours, minimum distance 2, Euclidean. Syllable type added as an axis for improved readability. Colours indicate specific individuals. See the full interactive figure in online Supplementary Materials, https://osf.io/jwt3g/?view_only=a971401f45744a709bb8accd0ee9769f

All parameters had PIC values greater than 1, indicating their significance in identity coding. The significantly highest PIC was observed in the coefficient of variation of IOI (i.e., how variable the IOIs were for an individual; Table 3).

**Table 3.** Potential of Identity Coding of the nine temporal parameters.

| Parameter | PIC value |
| --- | --- |
| Mean syllable duration (s) | 1.65 |
| Mean IOI (s) | 1.70 |
| CV syllable | 1.57 |
| CV IOI | 5.87 |
| Duration (s) | 1.37 |
| No. syllables | 1.54 |
| Syllable rate (Hz) | 1.75 |
| Rk global min | 1.32 |
| Rk global max | 1.40 |

### Does dissimilarity decrease with distance

Overall, vocal dissimilarities were highest for neighbours within 50 metres (Figure 5). We observed a negative statistical correlation between the dissimilarity score and the distance between singers in the mid-distance groups (a tendency to decrease up to 300 metres and a significant relationship up to 450 metres), but not between the closest or most distant singers (Table 4).

**Table 4.** Kendall’s coefficient scores for eight distance groups representing the maximum distance between the neighbours in a given group.

| Distance [m] | *p*-value | tau |
| --- | --- | --- |
| ≤50 | 0.691 | 0.149 |
| ≤150 | 0.874 | -0.022 |
| ≤300 | 0.072. | -0.133 |
| ≤450 | 0.012* | -0.126 |
| ≤600 | 0.009** | -0.115 |
| ≤750 | 0.557 | -0.023 |
| ≤900 | 0.749 | -0.032 |
| ≤2000 | 0.952 | 0.002 |

**Figure 5:**
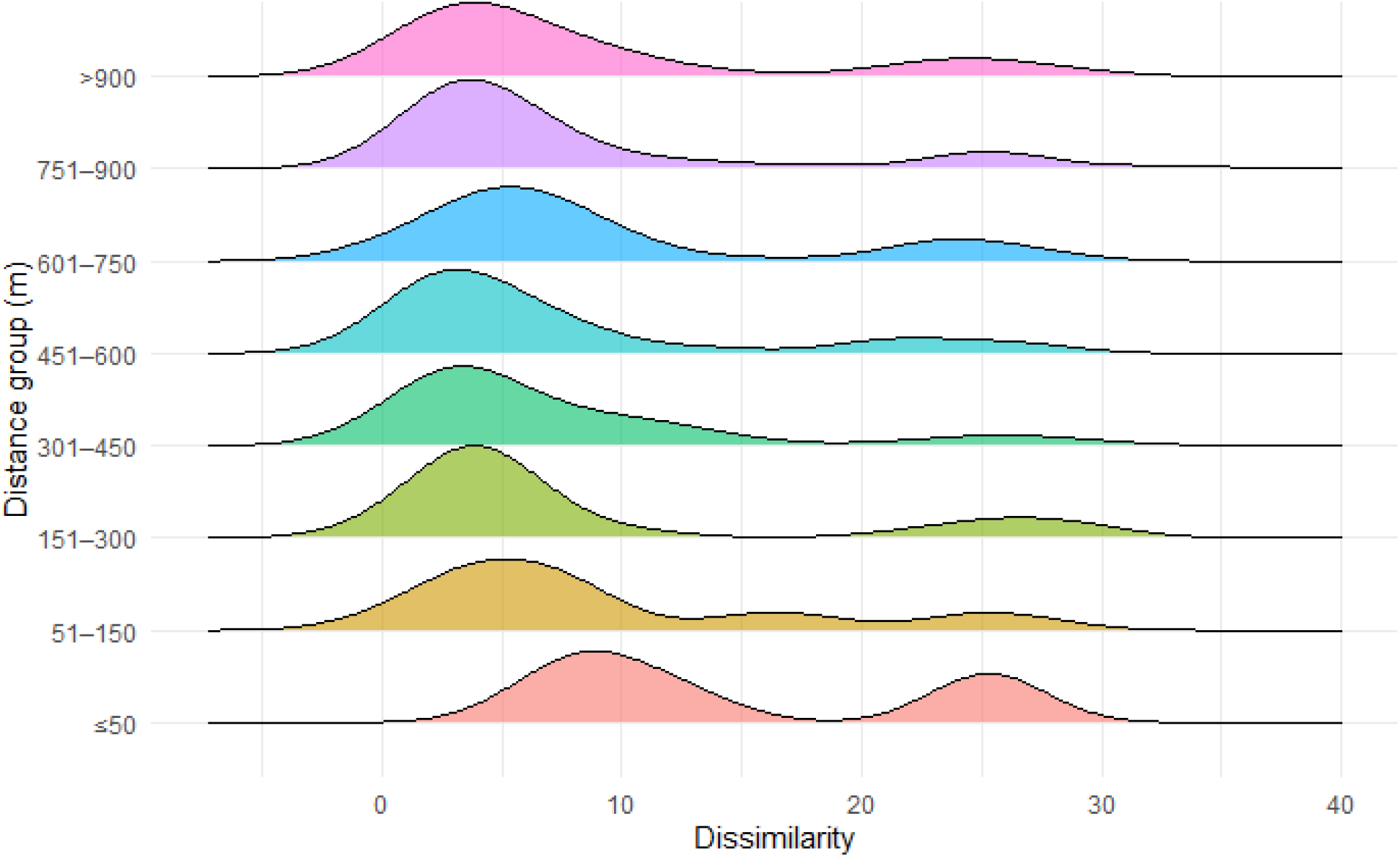
Density plots of dissimilarity scores for eight distance groups.

## Discussion

Using multiple songs of known individuals, we showed that the rhythmic parameters of the isochronous songs sung by yellowhammer males carry sufficient information to successfully assign them to a singer within and across phrase types. Rhythmic dissimilarities between two singers were the largest between the closest neighbours and decreased with the distance between them, but only at limited range and not over the entire locality, indicating that the singers may adjust rhythms to differentiate from their nearest neighbours.

Individual recognition has previously been suggested as one of the potential functions of isochrony Grabner et al. (2025). Amongst other features, certain basic temporal parameters such as tempo or coefficient of variation of syllable duration and inter-onset-intervals have been shown to carry individual variation in non-song displays of the Northern elephant seals *Mirounga angustirostris* Mathevon et al. (2017) and some seabirds Osiecka et al. (2024) Robisson (1992). To our knowledge, this is the first work to indicate that rhythms alone can inform on the passerine singer. In yellowhammers, the coefficient of variation of inter-onset-intervals (IOIs) within isochronous sequences and the syllable rates, as in syllables per second (to a lesser extent) showed by far the biggest potential of identity coding (PIC). This shows that within a predictable sequence of an isochronous song, small deviations can carry important information. The second most important parameter, syllable rates, has been shown to correlate with the arousal of the caller in some other species Filippi et al. (2019), which could influence our results. While we could not control for the singers’ arousal, all calls used in this study were recorded during solo territorial displays by adult males, therefore we expect arousal levels to be comparable between individuals. Another aspect is that deviations from isochrony tend to make a sequence more salient and easier to notice Van den Broek & Todd (2003). The variability of IOIs thus seems to emerge as the most important rhythmic parameter for coding the identity information. While other rhythmic parameters, such as e.g. beat precision Burchardt et al. (2021) might be of interest here, we have decided against using them (1) to avoid oversaturating the data with too many parameters, and (2) because the beat precision algorithm had been undergoing improvements during the preparation of this manuscript. However, we would like to highlight that the choice of rhythmic parameters should always consider both the current best practices and the characteristics of the studied sequence.

Interestingly, individual temporal patterns could be relied on both within and across phrase types, that is to say, across different songs. In real life, the situation is, of course, much more complicated. Birdsong is one of the most complex vocal displays known to science, and heavily influenced by culture Garland & McGregor (2020), singer’s investment in vocal exercise Adam et al. (2023), predisposed motor biases Toji et al. (2024) and choices of which parameters to focus on Goller (2022), among others. Rhythms do not exist in a vacuum, but rather strengthen the cues carried by the spectral parameters of a multidimensional acoustic signal and the spatial cues from the singer’s whereabouts. This study should thus be seen as a “dissected” investigation of one dimension of sound. The fact that the correct assignment to the individual was better across than within phrase types shows that these phrase types have, as expected, a rhythmic structure to them. We suggest that future efforts aiming to describe individuality in vocal sequences (such as e.g., the development of machine learning methods for population monitoring using passive acoustics) should use an enriched approach, combining both spectral descriptions and rhythmic patterns, and/or including training the image-based networks for rhythmic structure detection. Automating the process would also make this approach available to passive acoustic monitoring censuses that use acoustic capture-recapture for individual recognition.

While isochronous rhythms are used for purposes such as attracting female attention Norton & Scharff (2016) and facilitating duetting Rę k & Magrath (2023), the extent to which birds perceive rhythmic parameters is still poorly understood, and is likely to vary across species ten Cate et al. (2016). In birds that are not vocal learners, rhythms can be genetically determined and used as indicators of their evolutionary history Garcia et al. (2020) Sebastianelli et al. (2024). On the other hand, in open-ended learners, birdsong is often culturally transmitted and new songs and patterns can be learnt over the singer’s lifetime Garland & McGregor (2020). Yellowhammers, however, are close-ended learners, i.e. maintain their song type repertoires after the first year of their life. Overall, rhythm is the aspect that presents the biggest potential for cognitive control within a vocal display James et al. (preprint), allowing singers to adjust their output to a large extent. But why might yellowhammer neighbours come to sing particularly different rhythms? Matching or diverging from another’s calls is thought to carry intention or meaning in birds. For example, orange-crested conures (Eupsittula canicularis) respond more to calls that resemble their own Balsby et al. (2012), and matching/divergence has been suggested to serve as a means of pre-interaction negotiation in the species Balsby & Bradbury (2009) and signalling affiliative/antagonistic intentions Bradbury & Balsby (2016). For songbirds, who use their songs in sexual and territorial displays, this divergence may similarly not only maximise the chances of being distinguished from a neighbouring rival, but also serve as a message to such rival. Considering birdsong as an antagonistic display Searcy & Beecher (2009), stable rhythmic diversification from the neighbours observed in our study seems to follow the same pattern.

Social impacts on vocal diversity (e.g. Smeele et al. (2025) Madhavan et al. (2025)) and similarity (e.g. Luef et al. (2017) Baciadonna et al. (2022) Osiecka et al. (2023)) have been previously described in parrots, corvids, owls, and seabirds. Here, we add to this by showing that not only the social bonds, but also the spatial relationship between individuals, may contribute to their vocal diversity. Additionally, we suggest that the question of vocal individuality in general might be better approached at smaller spatial or behaviourally useful ranges. It seems plausible that one cannot sound very different from an entire population of conspecifics - but a diversification from neighbours or competitors can be possible and useful. This is somewhat similar to how a caller might need to sound particularly individual in groups of higher, but not lower, densities Madhavan et al. (2025).

The fact that we found no simple linear relationship to singer dissimilarity with distance (i.e., that the dissimilarity did not decrease linearly over the distance of two kilometers) is not surprising. The distance at which a pattern was observed (i.e. up to around 600 m) probably more than covers the active space of the species’ song, and it can be expected that there is no need to differentiate oneself from far away singers one does not come in interaction with. For comparison, the active space of a Red-winged blackbird (*Agelaius phoeniceus*, a North American songbird, was estimated at roughly 182 m in the absence of any wind Brenowitz (1982), and the real-life zebra finch (*Taeniopygia guttata castanotis*) song only travels up to roughly 9 m in nature Loning et al. (2022). Individual yellowhammer males establish territories of up to approx. 150 metres in diagonal Møller (1990), where they tend to stay in the same, preferred singing location, and we are confident that the observed pattern falls well within the useful ranges of the species. This work highlights the importance of rhythmic patterns in songbirds and suggests social impacts on rhythmic output in at least some vocal learners.

## Authors contributions

ANO: Conceptualization, methodology, formal analysis, investigation, data curation, writing: original draft, review and editing, software, visualisation, project administration, funding. JK: Investigation: data collection and processing. LSB: Conceptualization, methodology, formal analysis, writing: original draft, review and editing, software. MQS: Investigation: data processing, writing: review and editing. TP: Conceptualization, methodology, investigation: data collection, data curation, writing: original draft, review and editing, funding.

